# Pathway-Centric Visualization of Cell-Cell Communication in Single-Cell Transcriptomics Data

**DOI:** 10.1101/2025.07.25.666732

**Authors:** Sidrah Maryam, Martin Manolov, Rafael Kramann, Sikander Hayat

## Abstract

**Objective:** Single-nuclei transcriptomics enables investigating ligand-receptor mediated cell-cell communication between different cell-types. However, current tools do not allow for non-programmatic means to access this analysis. Additionally, most methods can not account for molecular pathways involved in downstream receptor signaling in cell-cell communication.

**Methods:** We developed scVizComm, a ShinyApp based web-portal that can be used to interactively visualize ligand-receptor networks across cell-types and different conditions.

**Results:** scVizComm can be used to host pathway centric pre-calculated cell-cell communication results. Using human kidney and heart single-nuclei transcriptomics data, we showcase the utility of scVizComm in understanding ligand-receptor interactions involved in fibrosis related pathways.

**Conclusion:** scVizComm allows interactive visualisation and pathway centric prioritisation of cell-cell communication analyses.

## 1 Introduction

Computational tools to analyze cell-cell communication (CCC) have become a mainstay of typical single-nuclei transcriptomics (snRNA) analyses. CCC is pivotal to understanding cellular crosstalk that governs cellular behavior in homeostasis, differentiation and disease progression. Multiple computational tools have been developed to identify putative ligand-receptor interactions that mediate CCC across cell-types [1]. Moreover, network topology based methods have also been developed to identify condition-specific ligand-receptor interactions [2]. Recently, a few tools have also been implemented to infer downstream activity of potential ligand-receptor interactions[3]. First generation of CCC methods provided a ranked list of condition agnostic ligand-receptors, while methods like CrossTalkeR spearheaded the comparative analyses across conditions [4]. Some methods can also compare CCC across samples [5]. However, all these methods require programmatic handling which severely limits the use of these tools for non-coders. Additionally, most tools do not account for underlying well-studied genesets and pathways that might play a role in downstream activity due to the putative ligand-receptor interactions.

To overcome these limitations, we have created scVizComm, an interactive visualization tool to display pathway and associated ligand-receptor interactions. This shiny app implementation of the tool would allow users to explore deeper insights of the cellular interactions obtained from fundamental CCC methods. scViZComm has four unique features 1) Visualise condition-wise Ligand-Receptor interaction for the source and target clusters of choice, 2) Determine expression dependent LR Score, 3) Distribution of genes associated with the selected pathway using AUCell [6] 4) KEGG pathway [7] analysis for the receptors associated per cluster or condition, thereby determining the downstream of the receptor. scVizComm can be accessed online through our shinyApp.

|  |  |
| --- | --- |
| Problem Issue | It is challenging to interactively visualise cell-cell interactions in scRNA data. |
| What is already known | Cellular interactions at the sample level and downstream signalling genes across diseases |
| What paper adds | The usefulness of the existing CCI tools is increased by including pathway-level data and permitting comparisons on cell type and disease level in an interactive, graphical program. |
| Who would benefit from the new knowledge in this paper | Biologists and clinician scientists can investigate CCI and ligand-receptor interactions on multiple levels in non-programmatic approach |

## 2 Materials and Methods

scVizComm relies on fully quality controlled (QC)-ed and annotated datasets. It requires two essential columns for cell type annotation and sample condition (e.g. disease, perturbation, treatment). Cell type annotation present in the column name ‘cell type original’, and condition or groups in ‘treatment’. The entire framework written in R language consists of several steps - i) Inferring L-R interaction, ii) determining AUCell ranking of pathway, iii) calculating differentially expressed genes in the data using MAST [8], and iv) exploring the CCC output results in scVizzComm.

The first step involves computational processing of the data.

### 2.1 Step 1

#### 2.1.1 Determining ligand-receptor interactions

Once the data is annotated and categorised into different cell types and conditions, the seurat object is converted to SingleCellExperiment(1.26.0) object and the counts matrix is extracted for each condition [9]. Then liana wrap function using ‘cell-phonedb’ as the method parameter is used to infer ligand receptor relationship for each interacting cluster for every condition using liana(0.1.13) [10].

#### 2.1.2 Determining AUCell Score

The AUCell run function from AUCell(1.26.0) package is employed to determine the scores for each cell utilising Canonical C2-KEGG pathways for pathway information[6].

#### 2.1.3 Differentially Expressed Genes

The differentially expressed genes are computed using FindMarkerGenes function in Seurat, implementing MAST method(1.30.0) [8]. The parameter logFCthreshold value was set to 0 and only.pos was set to False to report all differential genes with all fold-change values.

#### 2.1.4 Determining ligand-receptor(LR) Score

The interpretation of ligand-receptor interactions in step 1 provides the ligand-receptor information along with the source and target cluster. The ligand.expr and receptor.expr columns in the data frame obtained as an output of the cell interaction analysis provides the expression of ligand and receptor in the source and target clusters respectively. To quantify the strength of ligand-receptor interactions, an LR score was computed by applying the geometric mean to the ligand and receptor expression values. This approach ensures that higher expression of ligand and receptor results in elevated scores, thus penalizing imbalance pairs where one partner is weakly expressed, thereby, reflecting better biological scenarios where both components are essential for effective signaling.

The formula used for calculating the score:

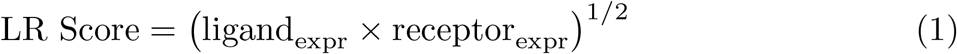

The pre-processing of the data can be performed using scVizzComm framework, and the resulting output files serve as the input for shiny app. The application offers a range of interactive, customizable features to facilitate in-depth exploration and interpretation of the cell-cell communication landscape derived from the analysis outputs.

### 2.2 Step 2

#### 2.2.1 Dataset Overview

scVizzComm can be utilised to schematically conduct basic data exploration on a 2D UMAP embedding to display gene-expression, and the number of cells in each cluster and condition. The users can provide any gene of interest to check its expression in each cluster and condition. Additionally, the app provides heatmap and feature plot to represent the expression in clusters and condition, and visualising the gene on data respectively.

#### 2.2.2 Circos Plots

Based on the LR Score calculated from CCC output (see section - Determining ligand-receptor(LR) Score), the overall interaction between different cell types is visualized. The edge thickness determines the number of ligand-receptor interactions and the weight determines the LR Score.

#### 2.2.3 Differential Circos Plots

Here, we present the differential expression changes in cellular communication between conditions of interest. The two conditions of choice are selected by the user and the Circos plot representing the differential LR Scores for interactions are plotted. The differential LR Scores are calculated using:

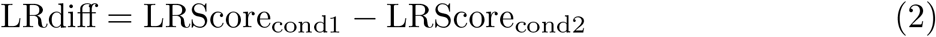

Where, LRScore_cond1_ and LRScore_cond2_ represent the LR scores for *condition1* and *condition2*, respectively. The LRdiff quantifies the extent to which *condition1* is perturbed relative to *condition2*.

#### 2.2.4 Condition-specific Ligand-Receptor Relationship

In order to investigate the ligand-receptor-pathway (LRP) relationship, one of the respective nodes (e.g. ligand-receptor-pathway) is chosen by the user. Once a node type is selected, the corresponding ligand/receptor/pathways are presented on the alluvial graph. An alluvial graph with edges pointing from ligands to receptors and then to pathway is created using the geom alluvium function from the ggplot2 package (3.5.1). Additionally, a receptor table is produced that depicts the number of ligands a receptor binds to in a given selected combination of source and target clusters, and number of pathways the receptor is linked to in msigdb database(7.5.1). The cell type specificity of a receptor is also determined by its presence in the differentially expressed genes for the target cluster.

#### 2.2.5 CCC-guided Receptor ranking and pathway scoring

Based on the user’s choice of the target cluster, all the receptors present in the cell type are ranked on the basis of LRmean score, where LRmean score is the mean of the LR score values for that receptor. Further, KEGG analysis is performed on all the receptors in the chosen target cluster and condition using clusterprofiler(4.12.6), that helps identify the downstream pathways associated with the receptors when they interact with various ligands [7]. On selecting one receptor from the target cluster, a list of pathways associated with the receptor is generated. To determine AUCell score distribution, once a pathway is selected from the list, we obtain the AUCell density plots in the entire data and condition. Besides, the AUCell score distribution is also obtained as boxplots in different cell types and conditions.

#### 2.2.6 Evaluation of differential Ligand-Receptor networks across conditions

Unique and common LR pairs can be determined on the basis of the user’s choice. Here, the user provides the conditions, source, and target clusters of interest, to establish the LR pairs. The scatter plot is produced where unique LR pairs for respective conditions are shown on the axis, while the common pairs are scattered based on the LR Score in the quadrant. Also, the heatmap showcases the fold-change difference between the conditions.

The perturbation between the two conditions is determined by the formula:

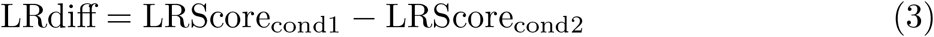

Where, LRScore_cond1_ and LRScore_cond2_ represent the LR scores for *condition1* and *condition2*, respectively. The LRdiff quantifies the extent to which *condition1* is perturbed relative to *condition2*.

#### 2.2.7 Condition-source specific KEGG pathway analysis

KEGG pathway enrichment analysis is performed for a selected source cluster under specific experimental conditions to identify significantly enriched pathways across multiple target cluster. The KEGG analysis is performed using the enrichKEGG function of package clusterprofiler(4.12.6)[7]. The receptors present in the target clusters that interact with the ligands from the source cluster, and thereby initiate the pathway activation are provided as an input to the KEGG analysis, and hence, determine the pathways that get activated on crosstalk.

#### 2.2.8 Source-Target specific condition-wise KEGG analysis

Here, for a given pair of source and target clusters, the KEGG pathway analysis is performed for every condition in the data using clusterprofiler(4.12.6) [7]. Consequently, for the same combination of interacting cell types, this aids in identifying the disease-specific pathway-level changes.

## 3 Results

We performed scVizzComm assisted data analysis on two different tissues - the human heart [11] and kidney [12] datasets as follows:

### 3.1 Use case 1: Interactive visualisation of ligand-receptors and downstream pathways in Myocardial Infarction (MI) Human data

Here, we used myocardial infarction data of human heart from left ventricle containing 191,795 cells to demonstrate the application of scVizzComm in visually exploring ligand-receptors and downstream pathways. Briefly, the data consists of four different conditions including control, tissue from the Fibrotic zone (FZ), myocardial infarction ischemic zone (IZ), and relatively unaffected area of myocardial infarction (Remote zone - RZ) [11]. We focused on significant ligand-receptor interactions obtained from cellphonedb, with p-value less than 0.05. In differential comparison analyses, we found overall decreased interactions in the fibrotic zone, however, most interactions were increased in the ischemic zone as compared to control (Figure 2A). For downstream workflow analyses, we focused on Cardiomyocytes interaction with mast cells in fibrotic zone with respect to control, to study the changes involving cell-cell communication between the immune and cardiac cells.

**Figure 1:**
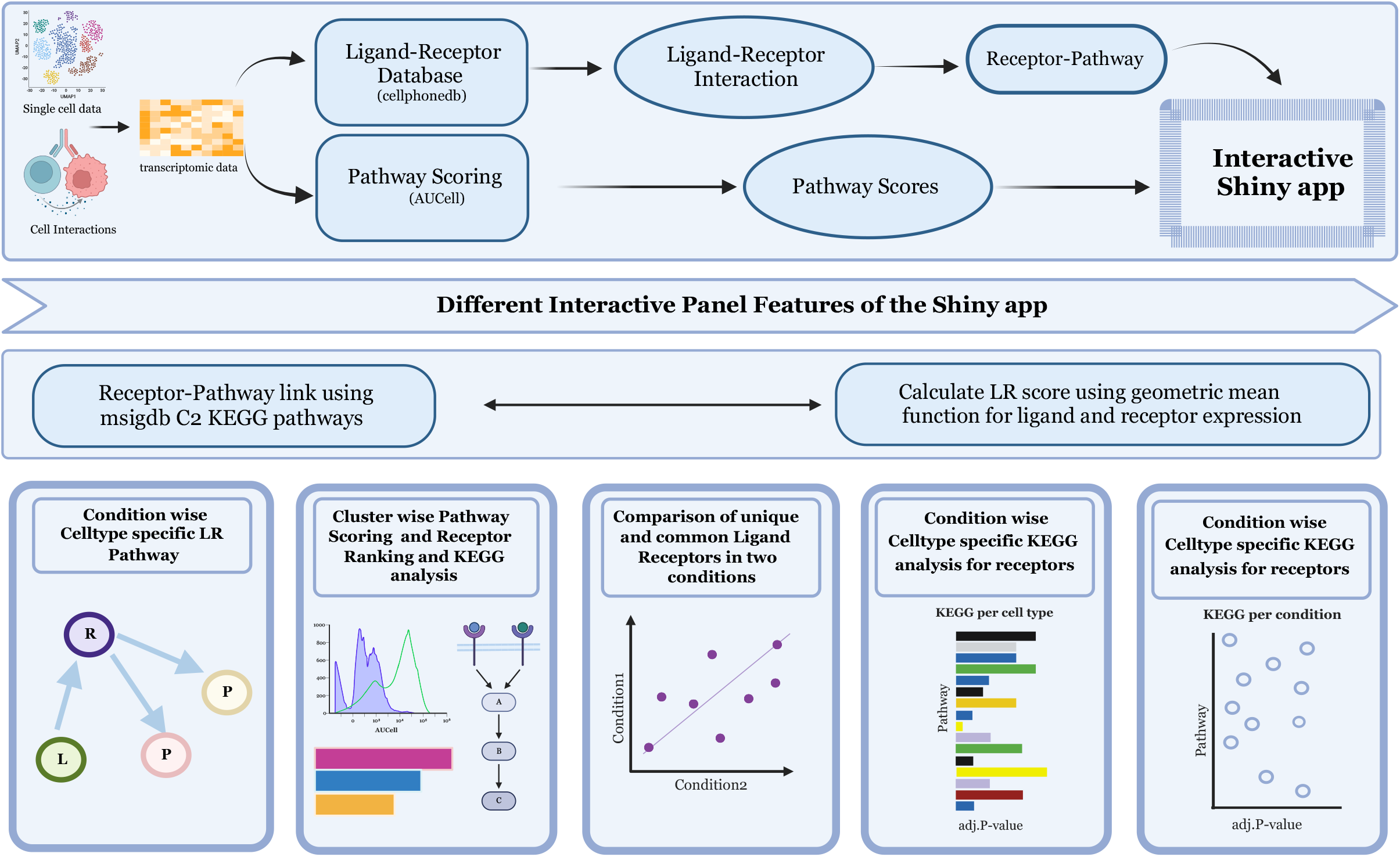
The schematic representation of the scVizzComm workflow. The preprocessing step is implemented on clustered and annotated data as a seurat object. Further, the ligand-receptor information is calculated using cellphonedb database, and AUCell is used for pathway scoring. The aggregated LR score is calculated using geometric mean. The ligand-receptor-pathway relationship is determined using msigdb. The output files from the preprocessing steps are used as input to scVizzComm for further exploration of cell-cell communication across conditions and cell-types.

**Figure 2:**
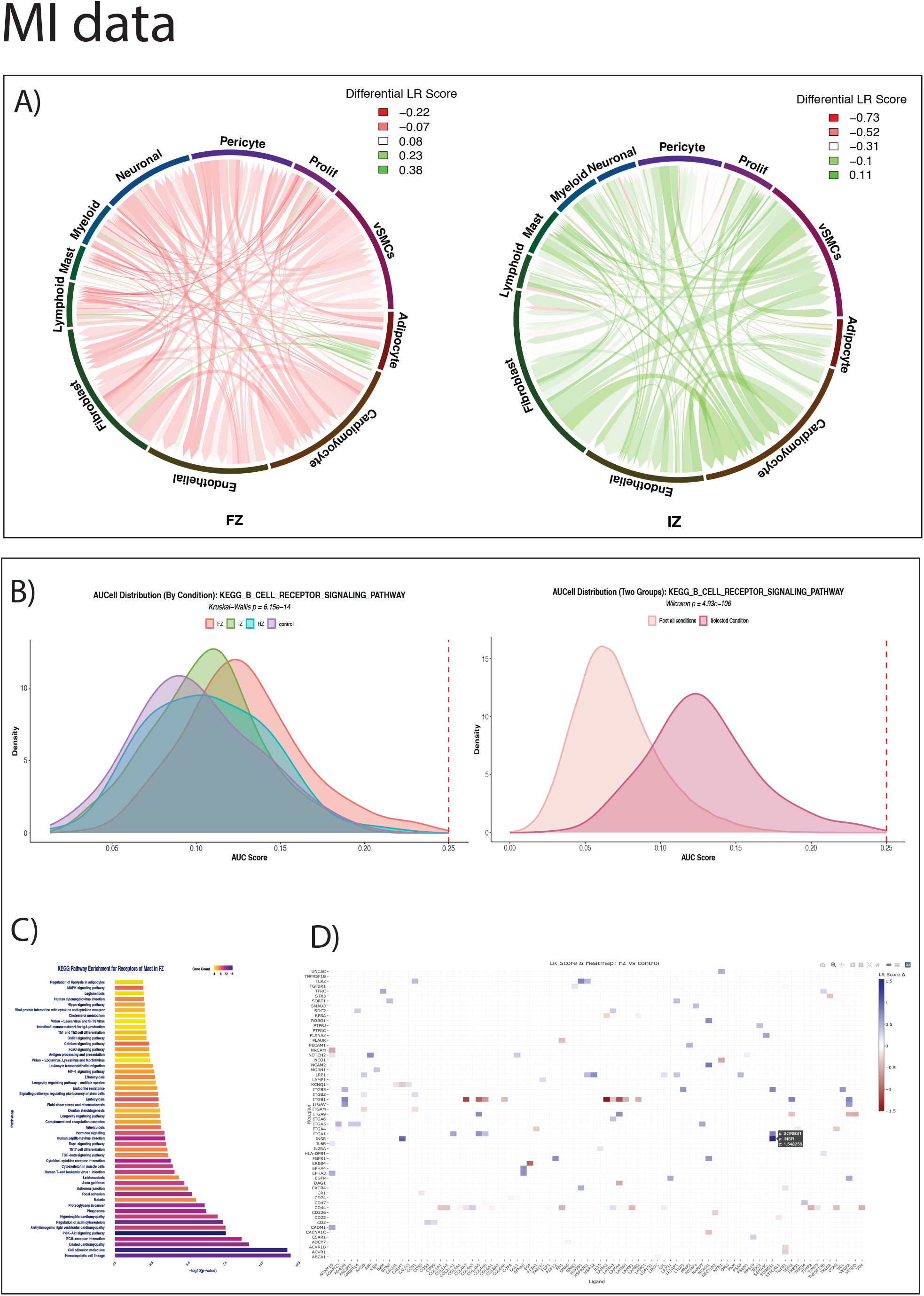
A) Comparative circos plots showing the LR score for change in cellular interaction in the 11 cell types for each FZ vs control, and IZ vs control. B) AUCell distribution plot for B cell receptor signaling pathway in FZ and other conditions. C) Bar plot depicting KEGG pathways associated with receptors expressed in mast cell clusters under fibrotic zone (FZ). The plot illustrates the number of overlapping genes within each pathway, highlighting an increase in pathway enrichment. D) Heatmap of selected ligand-receptor pairs and the change (LR_score FZ - LR_score control) in FZ vs control comparison for Cardiomyocyte as source and mast as target cell-type.

To showcase the ligand-receptor-pathway relationship, we selected NCAM1 as a representative receptor in FZ vs control comparison. NCAM1 (Neural Cell Adhesion Molecule 1) is a well known cell surface glycoprotein essential for mediating cellular and cell-matrix adhesion [13]. Additionally, NCAM1 is known to interact with multiple ligands and downstream effectors, positioning it as an ideal candidate to dissect potential pathway crosstalk in FZ condition. Increased expression of NCAM1 receptors were noted in the fibrotic zone, thereby suggesting the enhanced signaling in cardiomyopathy [14].

To further investigate the LRP relationship, we looked into the JAK-STAT signaling pathway, which is reported to be the inflammation pathway [15], was found to have more signal sending ligands in the fibrotic zone as compared to control(Figure 3). Next, JAK-STAT signaling pathway was chosen for in-depth analysis as it demonstrated a notable increase in signal-sending ligands within the fibrotic zone compared to control (Figure 3). Coupled with its established role in orchestrating inflammatory and fibrotic responses [15], this made JAK-STAT a biologically plausible and data-supported candidate for exploring ligand–receptor–pathway dynamics in our system.

**Figure 3:**
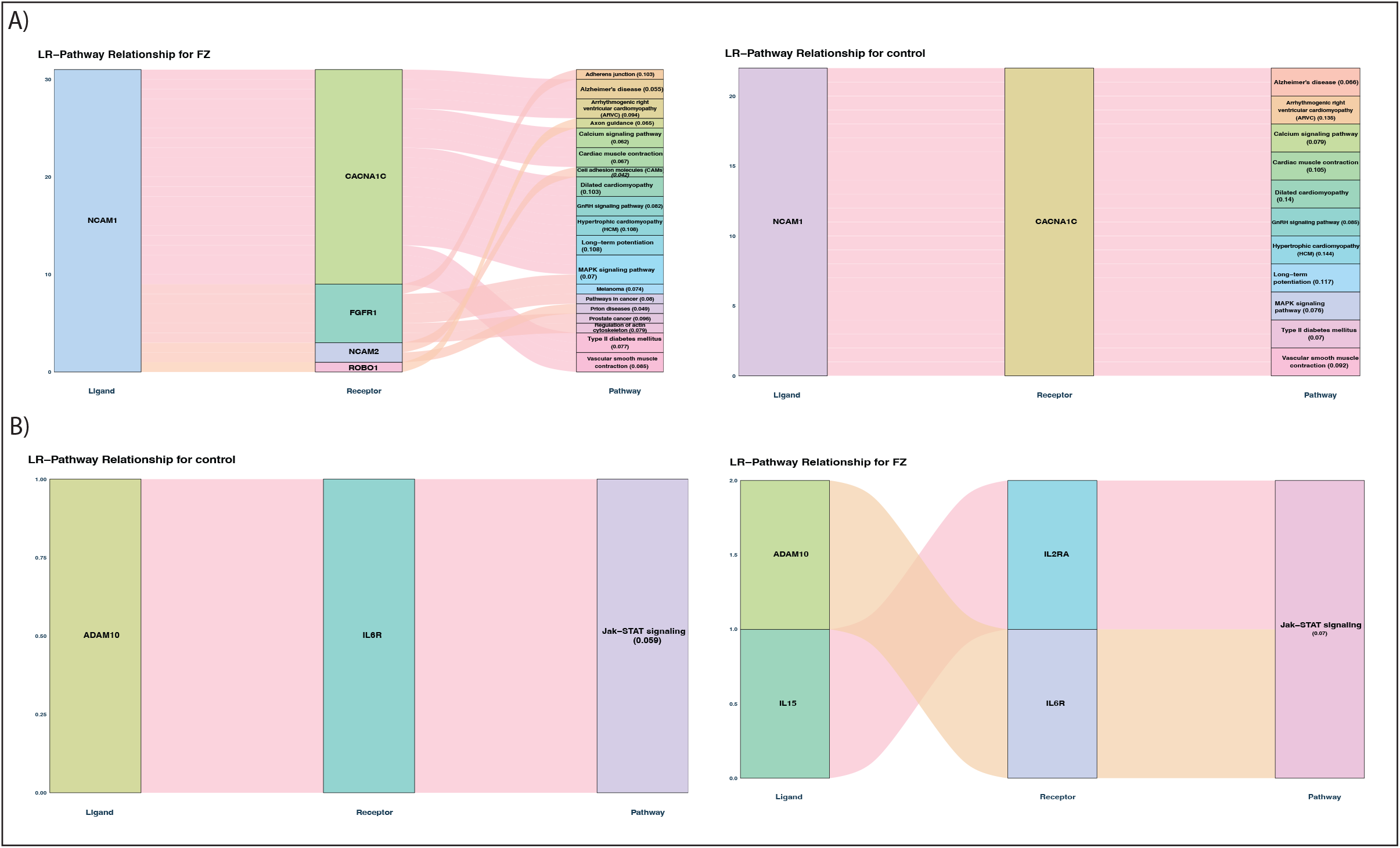
A) The alluvial graph showing the ligand-receptor-pathway relationship of selected ligands and pathways. B) NCAM1 ligand in mast cells in FZ (left) and controls (right) shows an increase in receptor interaction and overall change in downstream pathway activity. C) Pathway-centric analyses: For a selected pathway (JAK-STAT signaling node), all associated ligand and receptors in mast cells in FZ (left) and control (right) condition are shown.

Further, as proof of concept, we ranked the receptors present in the all celltypes using our LR score ranking, in each condition(Supplementary Table 1). We also ranked pathways for a selected condition and cell type where we chose mast cells as the target cluster(Supplementary Table 2). Next, the distribution of AUCell scores for genes associated with B-cell receptors was found to be significantly elevated in the fibrotic zone (FZ) compared to control tissue, consistent with previously reported immune activation in fibrotic cardiac microenvironment [16]. Additionally, KEGG pathway analysis of receptors expressed within mast cell clusters across various conditions revealed an enrichment of multiple immune-related signaling pathways, notably within ischemic and fibrotic zones. Pathways including Th1 and Th2 cell differentiation, Phagosome formation, and MAPK signaling exhibited notable up-regulation in these pathological contexts [15].

Furthermore, to identify condition-specific ligand-receptor interactions between cardiomyocytes and mast cells, a comparative analysis between the fibrotic zone and control tissue was performed. This analysis highlighted the SORBS1-INSR ligand-receptor pair as uniquely expressed within the fibrotic zone. Notably, this interaction has been previously implicated in the pathogenesis of hypertension [17], suggesting a potential mechanistic link within the cardiac niche (Figure 2B–D).

### 3.2 Use case 2: scVizzComm based analyses of the Kidney Precision Medicine Project (KPMP) data

The kidney dataset contain 200338 cells from Kidney Precision Medicine Project(KPMP) [12], consisting of healthy kidney tissue and several kidney diseases including chronic kidney disease(CKD), hypertension associated CKD (H CKD), diabetic kidney disease (DKD), acute kidney injury (AKI), and covid induced AKI (COV AKI). The differential patterns of cellular interactions observed in each pathological condition relative to the reference were found to be largely diminished (Figure 4A). To further investigate this phenomenon, we specifically examined the interactions between immune cells and proximal tubule cells, given their potential role in modulating local tissue responses.

**Figure 4:**
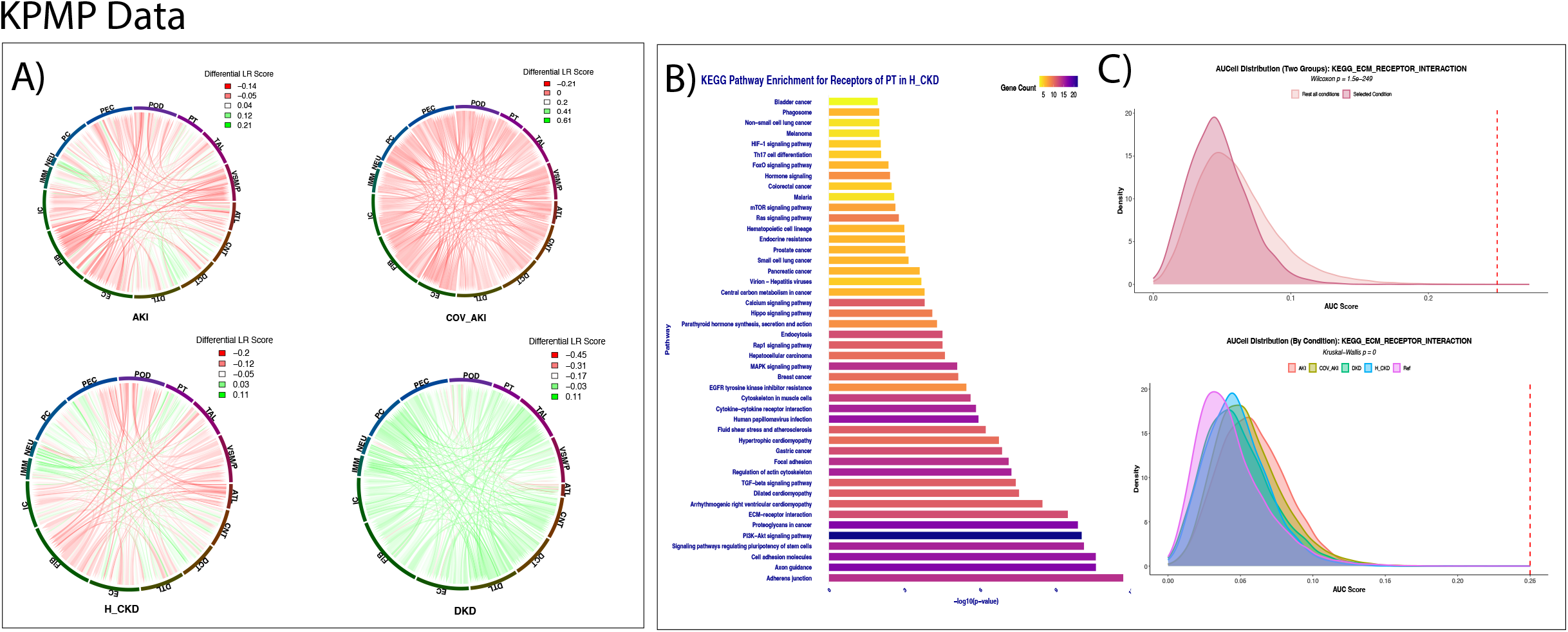
A) Circos plots illustrating cellular interaction networks across different cell types under various conditions, including acute kidney injury (AKI), hypertensive chronic kidney disease (H_CKD), diabetic kidney disease (DKD), and Covid-associated AKI, compared to the reference dataset. B) Bar plot depicting KEGG pathway enrichment analysis of receptors expressed within the proximal tubule (PT) cluster in control and H_CKD conditions. C) Distribution of AUCell scores for the extracellular matrix (ECM) receptor interaction pathway across individual conditions (top) and a comparative analysis of H_CKD versus all other conditions combined (bottom).

To determine the relationship between ligand–receptor–pathway interactions, we examined extracellular matrix (ECM)-related pathways in both the kidney injury and control conditions. Our analysis revealed heightened signal production and reception in the kidney injury group compared to the reference, consistent with the known association between excessive ECM signaling, kidney fibrosis, and functional disruption [18]. Given the pivotal role of the EGFR receptor in regulating cell proliferation, survival, and fibrotic remodeling, and its previously reported involvement in kidney disease progression [19], we selected EGFR as a representative node for further analysis. Notably, EGFR exhibited increased interaction density in the kidney injury condition (Figure 5A), underscoring its potential contribution to pathological signaling within the diseased microenvironment.

**Figure 5:**
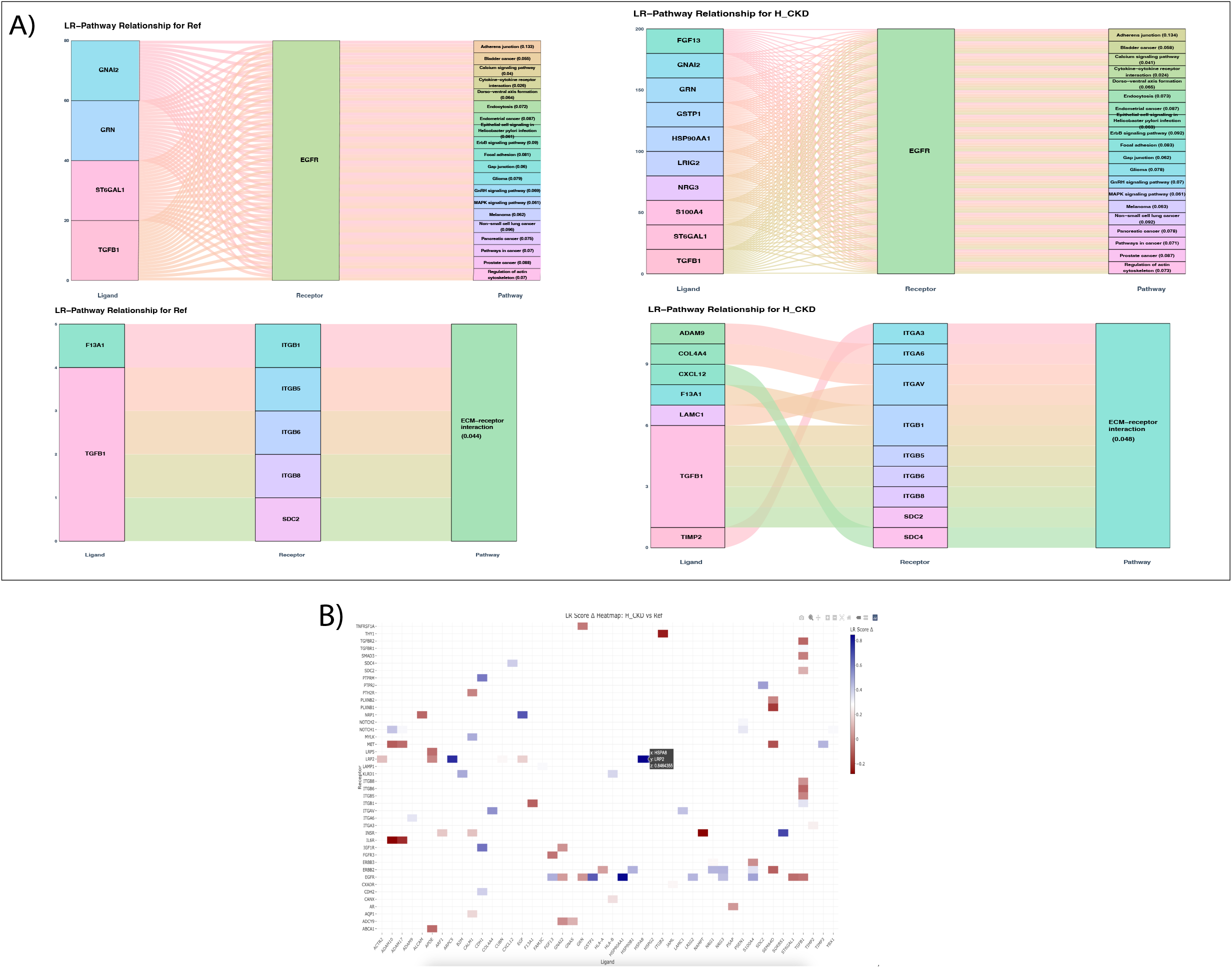
A) The alluvial graph showing the ligand-receptor-pathway relationship for selected ligand and pathway. Top - Here, we focused on EGFR receptor in Proximal Tubule cells and observed increased counts of ligands in H_CKD as compared to reference samples. B) Bottom - for ECM receptor interaction pathway, we found an increased number of ligands and receptors in the disease condition. Heatmap of selected ligand-receptor pairs and their change in LR_score as compared to reference samples.

Next, we ranked all receptors based on their ligand–receptor (LR) scores (Supplementary Table 3). We also ranked pathways associated with the Immune cells in H CKD(Supplementary Table 4).

To explore pathway-level changes in receptor expression, KEGG pathway analysis was performed on PT cell receptors. This revealed a notable loss of cholesterol metabolism pathways in the disease condition compared to the reference, consistent with previous reports linking altered lipid metabolism to kidney injury progression [20]. Additionally, we assessed the distribution of AUCell scores for genes associated with the extracellular matrix (ECM)–receptor interaction pathway across both reference and disease conditions. A higher AUCell score distribution was observed in the disease state, reflecting an upregulation of ECM-related signaling activity in kidney injury [21] (Figure 4B–C).

For the unique ligand–receptor pair analysis, we identified the HSP18–LRP2 pair as uniquely present and highly expressed in the H CKD condition, specifically within immune and proximal tubule cell interactions. The receptor has been previously implicated in mediating kidney injury and disease progression [22] (Figure 5B).

## 4 Discussion

snRNA-Seq allows us to address cell-cell communication which is mediated by ligand-receptor (LR) interactions and influences downstream signalling and pathways. While several computational tools exist for LR analysis, they often lack accessibility for non-programming users and typically do not incorporate pathway-level interpretation of downstream receptor signaling. To address these limitations, we developed scVizComm, a user-friendly ShinyApp that enables interactive exploration of precomputed LR networks across different cell types and biological conditions. One of the key innovations of scVizComm is its focus on pathway-centric analysis, allowing users to move beyond lists of LR pairs and towards understanding their functional roles within broader signaling cascades. Applying scVizComm to human kidney and heart snRNA-seq datasets, we demonstrate its utility in highlighting fibrosis-associated pathways, including TGF-*β*, Notch, and integrin signaling. This pathway-centric approach allows researchers to prioritize biologically meaningful LR interactions, particularly those converging on shared fibrotic mechanisms. Additionally, scVizComm facilitates hypothesis generation by providing an intuitive interface for exploring differential LR activity across conditions, such as healthy versus diseased tissue. Despite its strengths, scVizComm currently relies on precomputed LR inference results, meaning upstream data processing and statistical modeling must be performed externally. Future development could integrate these pipelines directly into the platform, allowing end-to-end analysis within a single interface. Nonetheless, scVizComm represents a significant step forward in making LR network analysis more accessible, interpretable, and functionally relevant. Its pathway-centric visualization paradigm offers a scalable and flexible solution for researchers seeking to uncover the mechanistic underpinnings of cell-cell communication in health and disease.

## 5 Conclusion

scVizzComm is a user-friendly shiny application that allows users to determine the downstream signaling of cellular interactions. It provides an accessible and interactive platform for visualizing ligand-receptor networks with a novel focus on pathway-centric interpretation. By enabling users to explore cell-cell communication in the context of biologically relevant signaling pathways, scVizComm bridges a critical gap between complex transcriptomic data and functional insight. Its user-friendly interface empowers researchers from diverse backgrounds to generate and prioritize hypotheses, particularly in disease contexts such as fibrosis. While future integration of upstream analytical pipelines will further enhance its utility, scVizComm already serves as a powerful tool for advancing mechanistic understanding of intercellular signaling in complex tissues. Importantly, the platform democratizes access to LR analyses, supporting scientists without extensive computational backgrounds and enabling collaborative interpretation in translational research settings.

## 6 Figure legends

### 6.1 Figure 1: Overview of the scVizzComm workflow

The schematic representation of the scVizzComm workflow. The preprocessing step is implemented on clustered and annotated data as a seurat object. Further, the ligand-receptor information is calculated using cellphonedb database, and AUCell is used for pathway scoring. The aggregated LR score is calculated using geometric mean. The ligand-receptor-pathway relationship is determined using msigdb. The output files from the preprocessing steps are used as input to scVizzComm for further exploration of cell-cell communication across conditions and cell-types.

### 6.2 Figure 2: Comparative analyses of human heart MI dataset

scVizzComm application on MI heart data. A) Comparative circos plots showing the LR score for change in cellular interaction in the 11 cell types for each FZ vs control, and IZ vs control. B) AUCell distribution plot for B cell receptor signaling pathway in FZ and other conditions. C) Bar plot depicting KEGG pathways associated with receptors expressed in mast cell clusters under fibrotic zone (FZ). The plot illustrates the number of overlapping genes within each pathway, highlighting an increase in pathway enrichment. D) Heatmap of selected ligand-receptor pairs and the change (LR score FZ - LR score control) in FZ vs control comparison for Cardiomyocyte as source and mast as target cell-type.

### 6.3 Figure 3: Ligand and pathway centric analyses of human MI data

The alluvial graph showing the ligand-receptor-pathway relationship of selected ligands and pathways. A) NCAM1 ligand in mast cells in FZ (left) and controls (right) shows an increase in receptor interaction and and overall change in downstream pathway activity. B) Pathway-centric analyses: For a selected pathway (JAK-STAT signaling node), all associated ligand and receptors in mast cells in FZ (left) and control (right) condition are shown.

### 6.4 Figure 4: Comparative analyses of the human kidney KPMP dataset

(A) Circos plots illustrating cellular interaction networks across different cell types under various conditions, including acute kidney injury (AKI), hypertensive chronic kidney disease (H CKD), diabetic kidney disease (DKD), and Covid-associated AKI, compared to the reference dataset. (B) Bar plot depicting KEGG pathway enrichment analysis of receptors expressed within the proximal tubule (PT) cluster in control and H CKD conditions. (C) Distribution of AUCell scores for the extracellular matrix (ECM) receptor interaction pathway across individual conditions (top) and a comparative analysis of H CKD versus all other conditions combined (bottom).

### 6.5 Figure 5: Ligand and pathway centric analyses of human kidney data

A)The alluvial graph showing the ligand-receptor-pathway relationship for selected ligand and pathway. Top - Here, we focused on EGFR receptor in Proximal Tubule cells and observed increased counts of ligands in H CKD as compared to reference samples. Bottom - for ECM receptor interaction pathway, we found an increased number of ligands and receptors in the disease condition. B) Heatmap of selected ligand-receptor pairs and their change in LR score as compared to reference samples.

## 7 Table

### 7.1 Supplementary Table 1

The ranked receptors from each cell type in each condition, based on LR score in MI data.

### 7.2 Supplementary Table 2

The ranked receptors from each cell type in each condition, based on LR score in KPMP data.

### 7.3 Supplementary Table 3

The pathway ranking for the pathways associated with receptors present in cardiomyocytes in FZ condition for MI data.

### 7.4 Supplementary Table 4

The pathway ranking for the pathways associated with receptors present in PT in H CKD for KPMP data.

### 7.5 Supplementary Table 5

Top 10 ligand-receptor pairs are listed for each celltype as source, for each condition in MI data.

### 7.6 Supplementary Table 6

Top 10 ligand-receptor pairs are listed for each celltype as source, for each condition in KPMP data.

## 8 Declarations

### 8.1 Ethics approval and consent to participate

The research was conducted ethically and in accordance with the relevant guidance.

### 8.2 Consent for publication

All authors approve for publication.

### 8.3 Availability of data and material

***Project Name*** : scVizzComm

***Project Home page*** : https://costalab.ukaachen.de/shiny/smaryam/

***GitHub:*** https://github.com/hayatlab/scVizComm

***Operating system:*** Platform independent

***Programming language:*** R

***Figures and Supplementary Tables:***

https://doi.org/10.5281/zenodo.15363559

### 8.6 Authors’ contributions

S.M. and S.H. conceived the experiment(s), S.M. conducted the experiment(s), S.M. analysed the results, M.M. supported the shiny App, S.M. and S.H. wrote and reviewed the manuscript, all authors edited and refined the manuscript.

