## Supplementary Figures for "Pathway-Centric Visualization of Cell-Cell Communication in Single-Cell Transcriptomics Data"

### Preprocessing data for application input

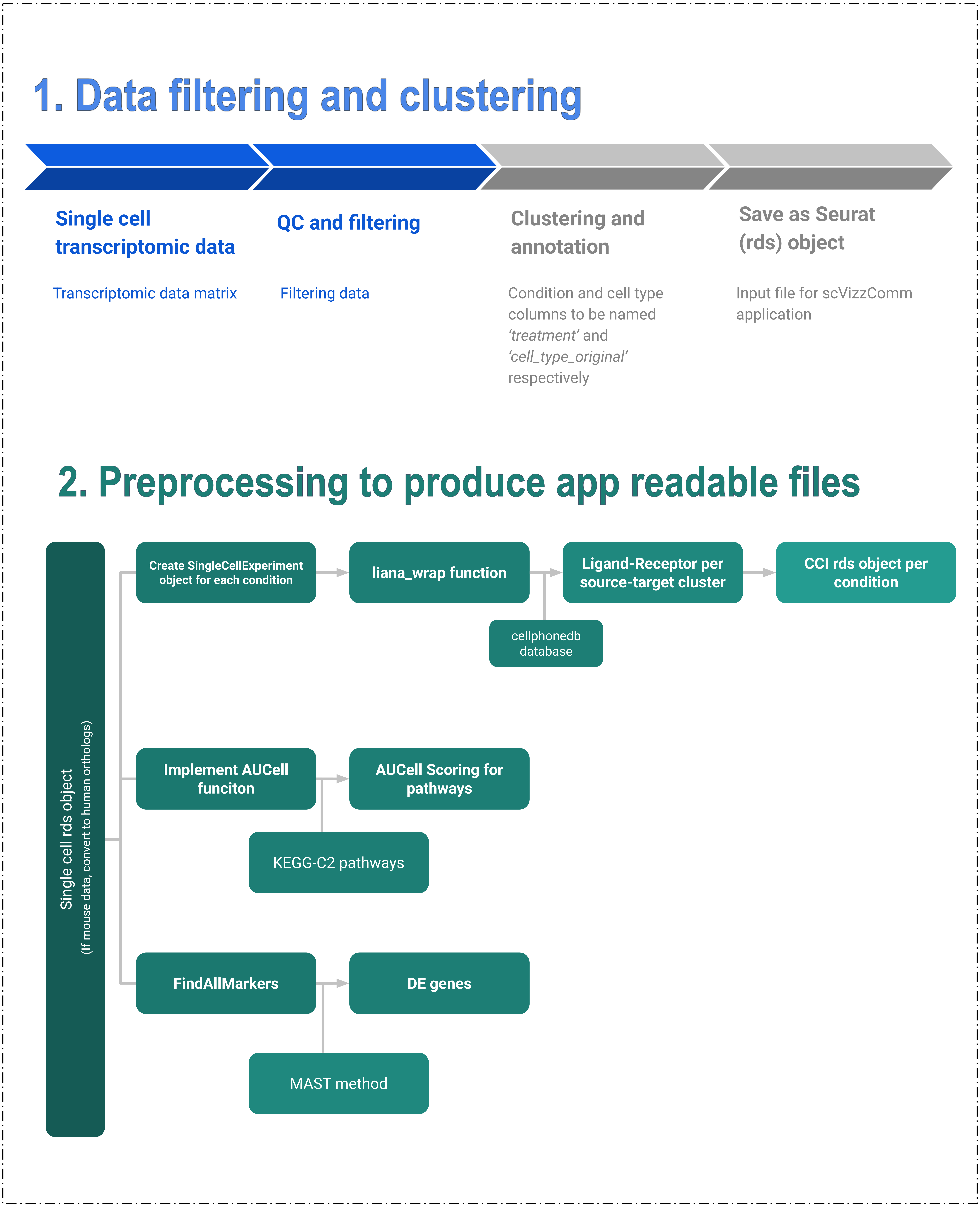

### 3. Modification of files by the app

#### Seurat Object

Required for condition and cell type information to plot figures and calculations

- None

#### CCI Files

To determine cellular interactions per cluster

- Only include significant interactions (pval <0.05)
- Map each receptor with curated msigdb KEGG-C2 pathways using omnipath
- Calculate LR Score using expression values of LR

#### AUCell score File

Determine expression of genes associated with specific pathway

- Subsetting condition wise and cluster wise pathway information using cell information
- Used KEGG-C2 pathways

#### DE Results

Determine the specificity of receptors

- Use pvalue < 0.05 for significance of receptors
- Presence of receptor in DE determines specific for particular cell type

### Panel - All Condition Circos Plots

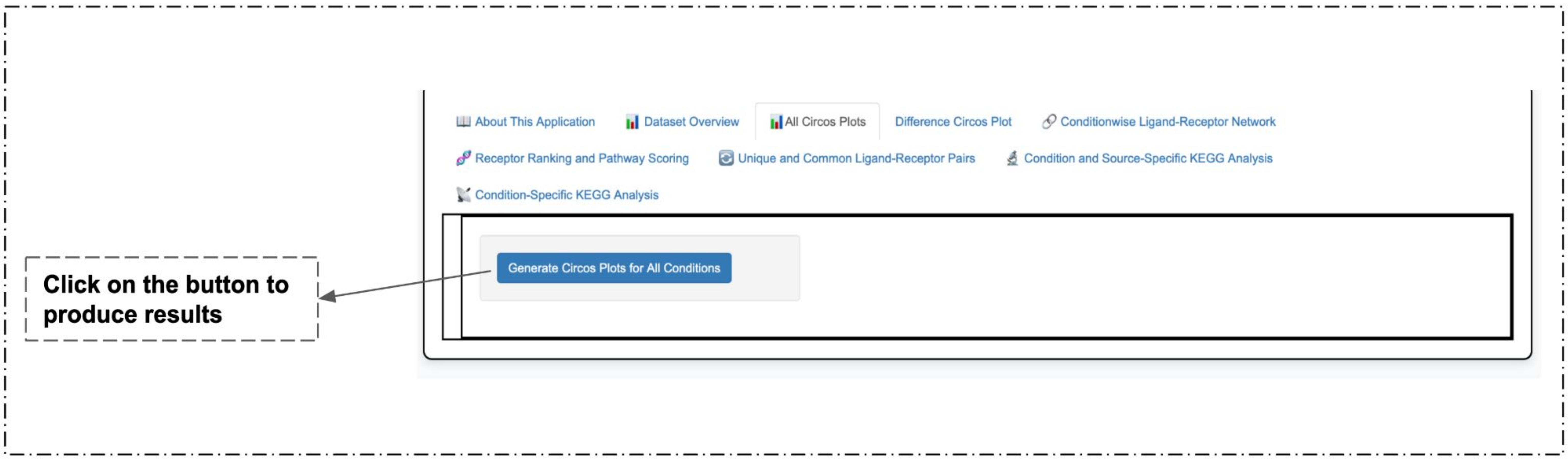


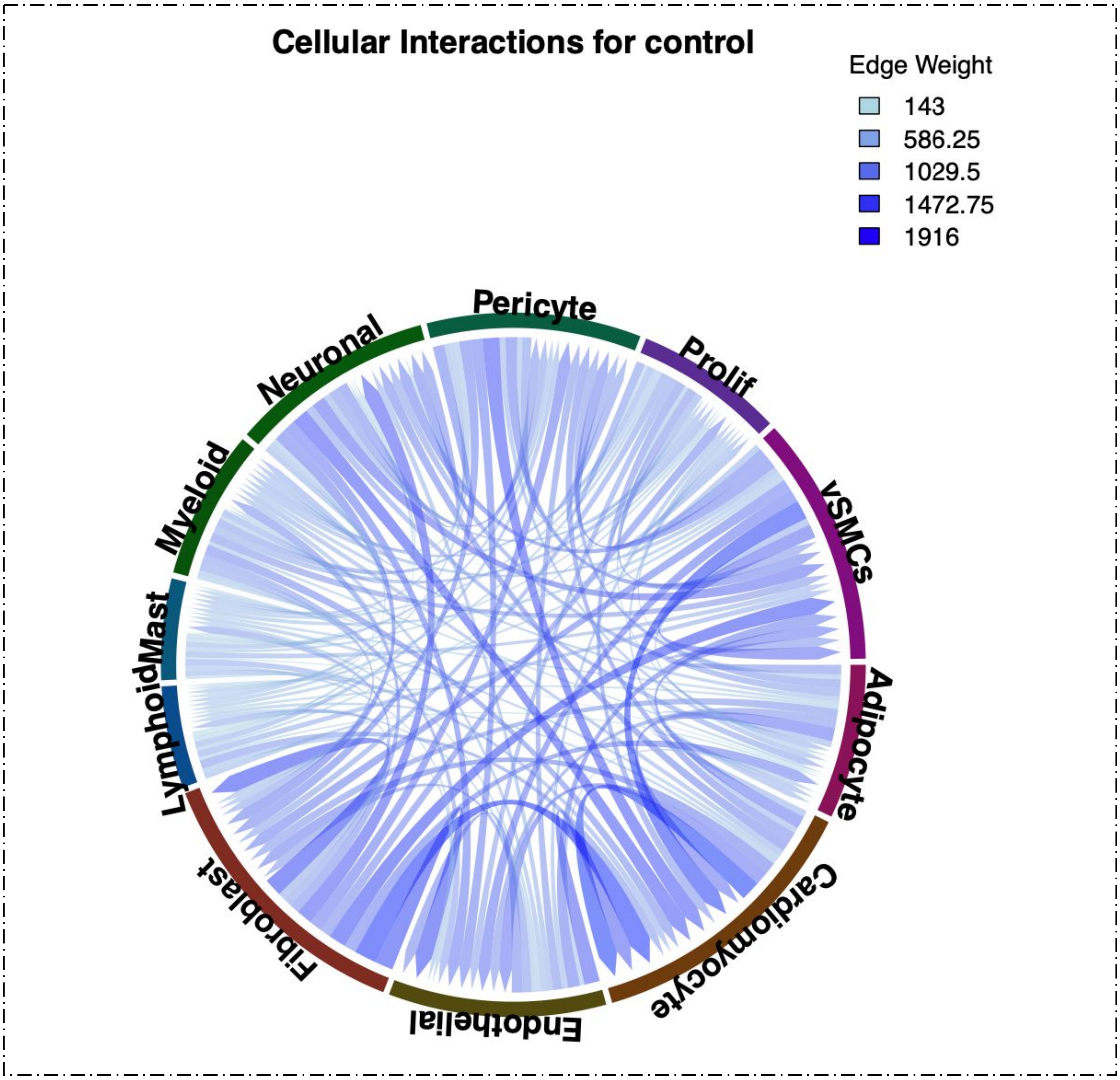


**Supplementary Figure 1.** Based on the Ligand-Receptor Score calculated using ligand and receptor expression obtained from cellphonedb analysis, the interactions are plotted per cell type for all the conditions individually.

### Panel - All Condition Circos Plots

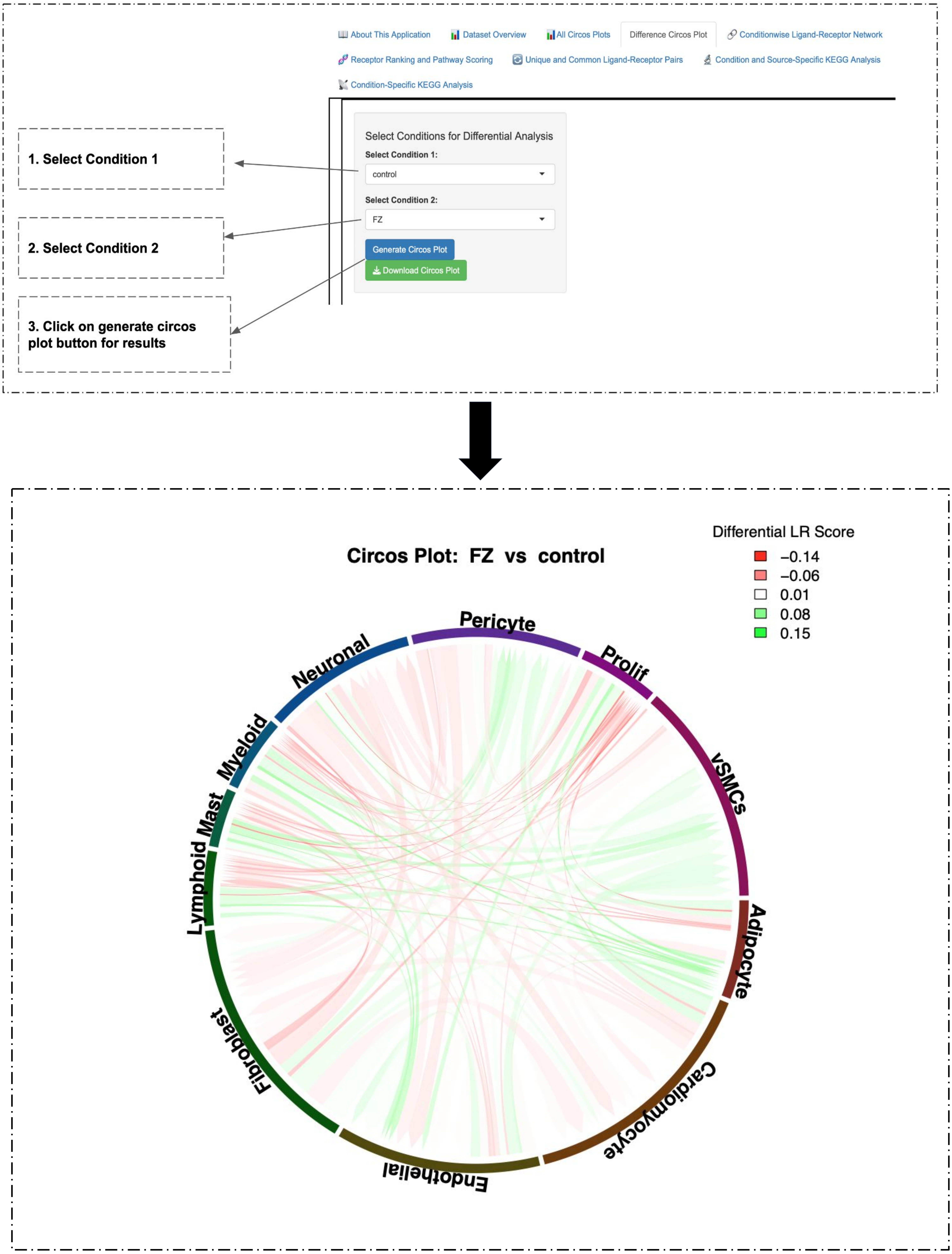


**Supplementary figure 3.** On choosing condition 1 and condition 2, and generating the figure, the perturbations in condition 1 with respect to condition 2 is produced. This is calculated using Condition 1 LR score - Condition 2 LR score. Red color denotes loss of interaction in condition1 wrt to condition2, and vice versa.

### Panel - Condition Wise Ligand-Receptor Network

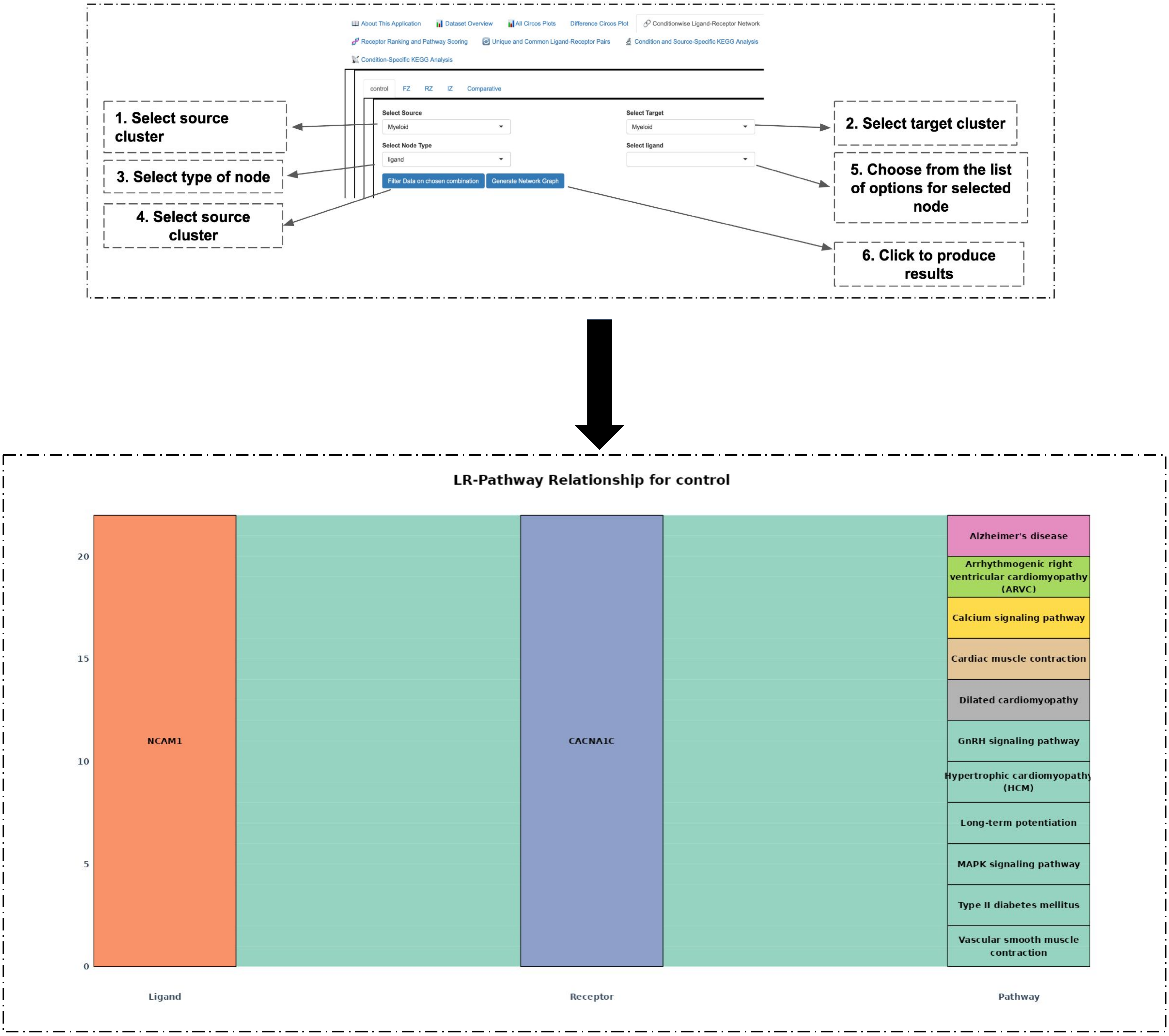


| Receptor | InDegree | NumberOfPathways | CellTypeSpecificity | Significance |
|---|---|---|---|---|
| CD74 | 1 | 1 | Yes | Significant |
| LRP1 | 1 | 1 | Yes | Significant |

**Supplementary Figure 3.** After following the instructed steps, ligand-receptor-pathway relation is obtained as alluvial plot, along with the table. The table gives the information about the receptor in the network, viz., number of ligands it binds to, number of pathways it is associated with and if it is cell type specific.

### Panel - Receptor Ranking and Pathway Scoring

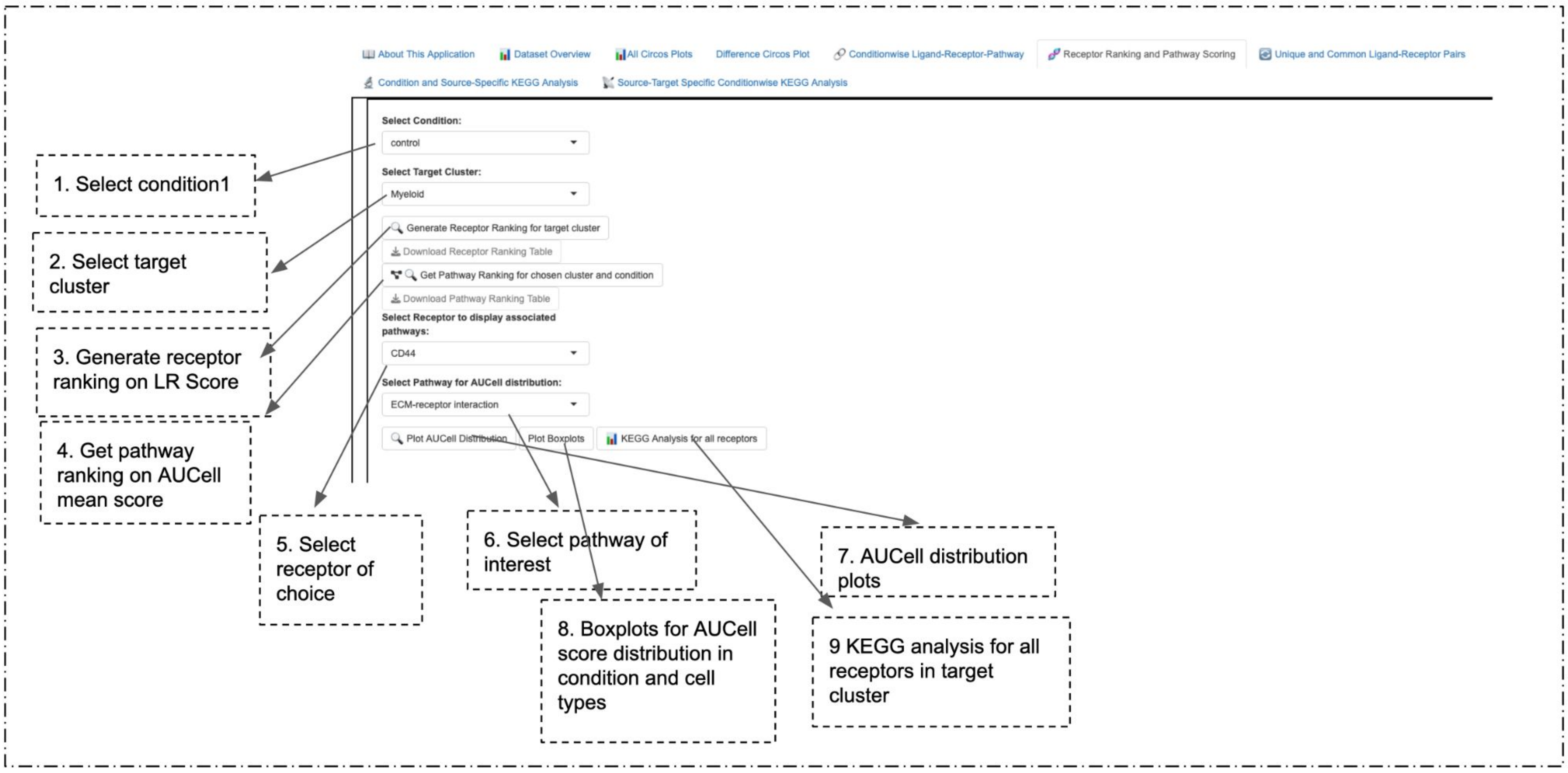


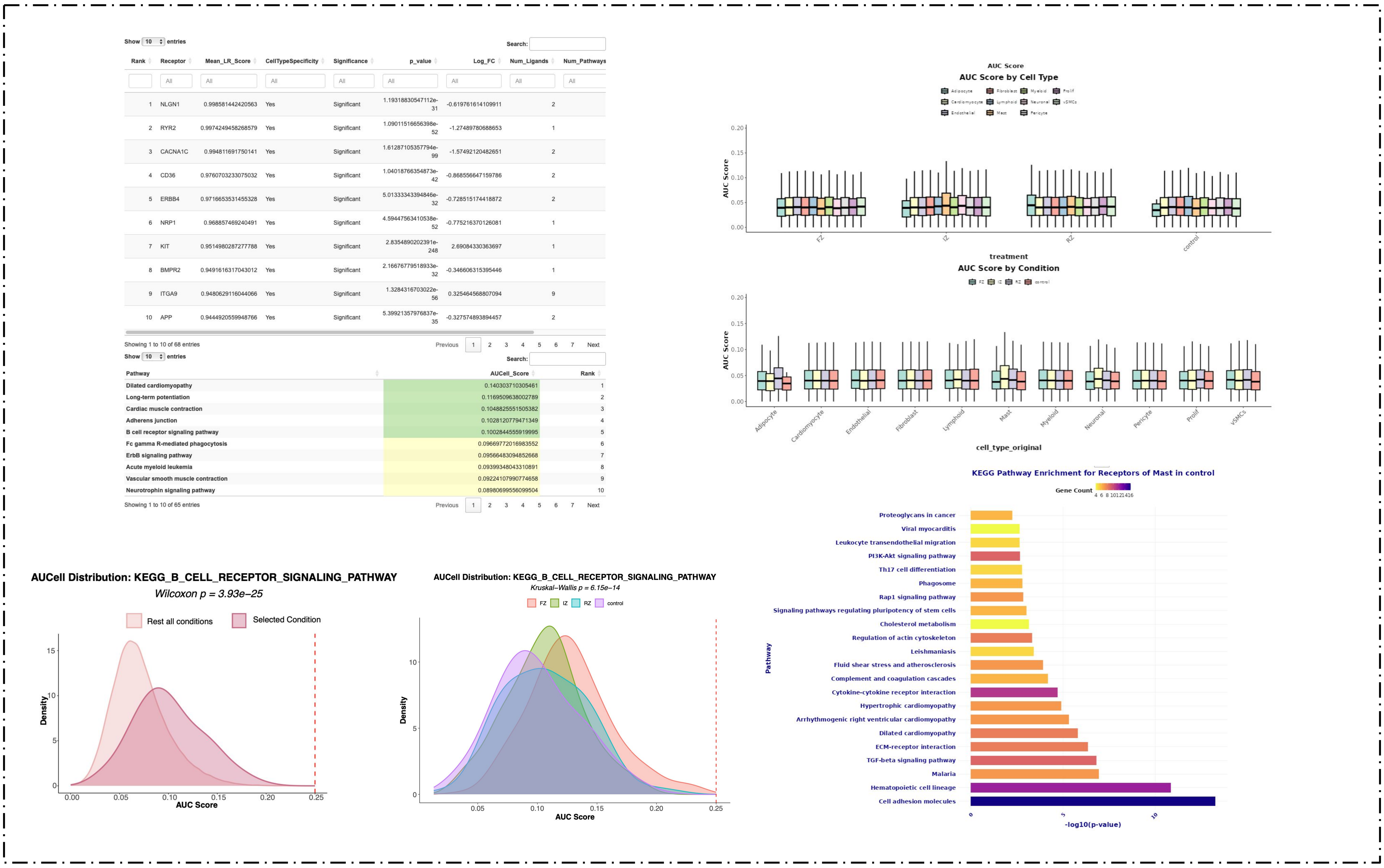


**Supplementary Figure 4.** In this panel, a condition and a target cluster is chosen, based on that the entire list of receptors present in the cluster, the receptor choice is listed. Also the ranking table for receptor is obtained on LR score. The barplot of enrichment KEGG analysis done on target-receptors is determined. On choosing receptor and associated pathway, AUCell distribution of the chosen pathway is produced.

### Panel - Unique and Common LR Pairs

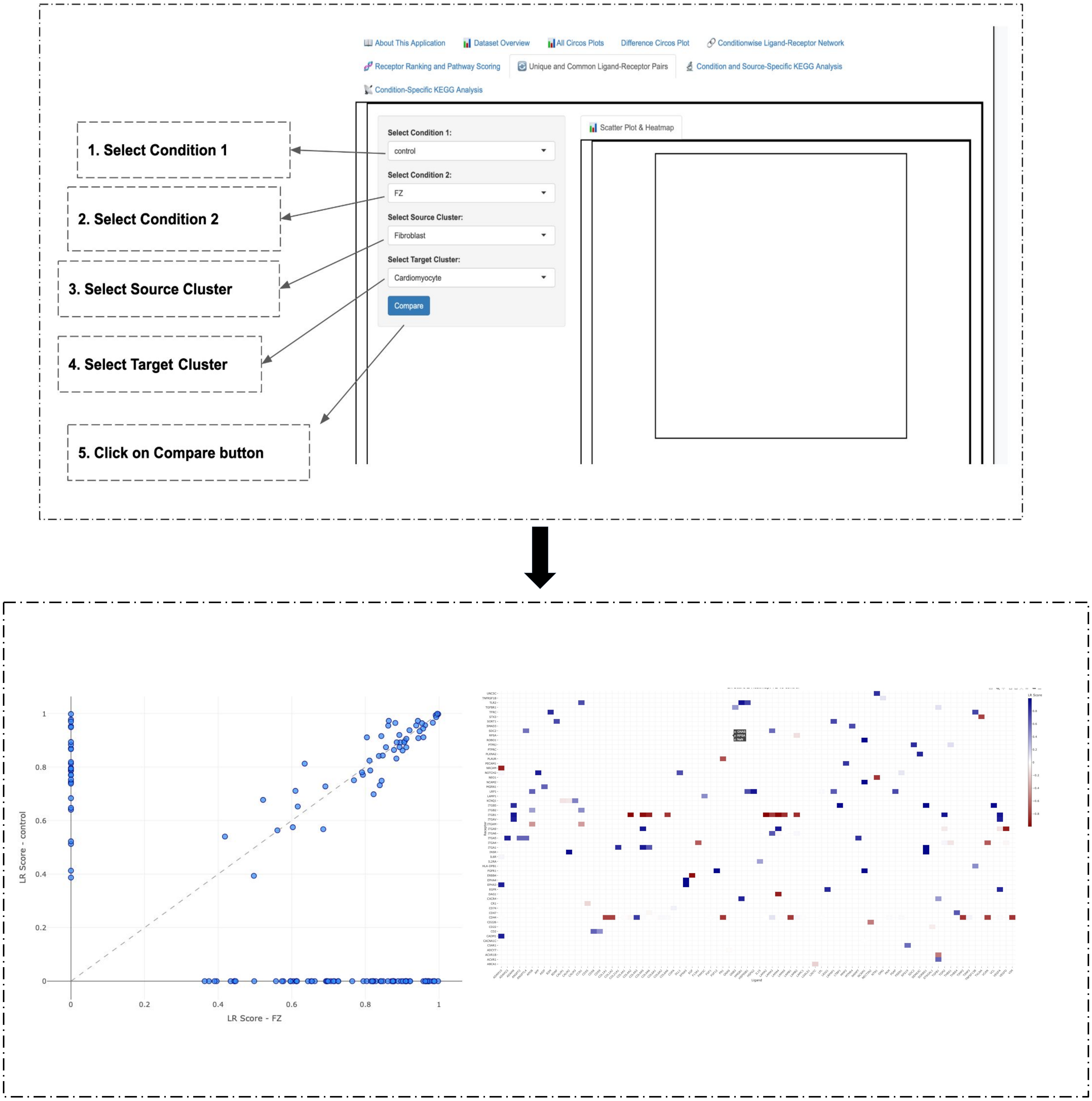


**Supplementary Figure 5.** On choosing condition 1 and condition 2, the ligand-receptor information for respective conditions are filtered on the basis of source and target clusters of choice. On generating the results, the unique LR pairs are plotted on x and y axis, while the common ones are scattered to show the deviation from each other. The heatmap shows the perturbation in condition 1 with respect to condition 2. The blue color denotes the higher LR score in condition 1.

### Panel - Condition and Source specific KEGG analysis

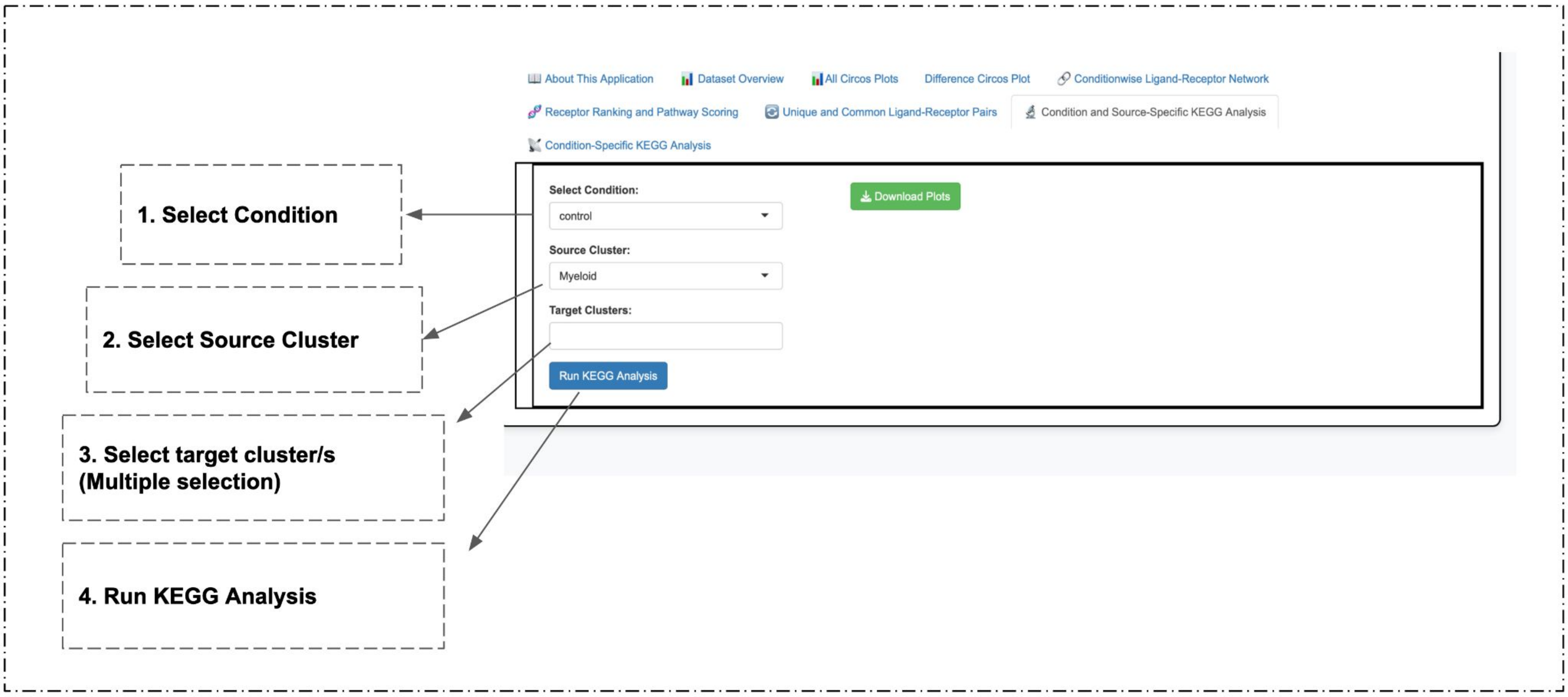


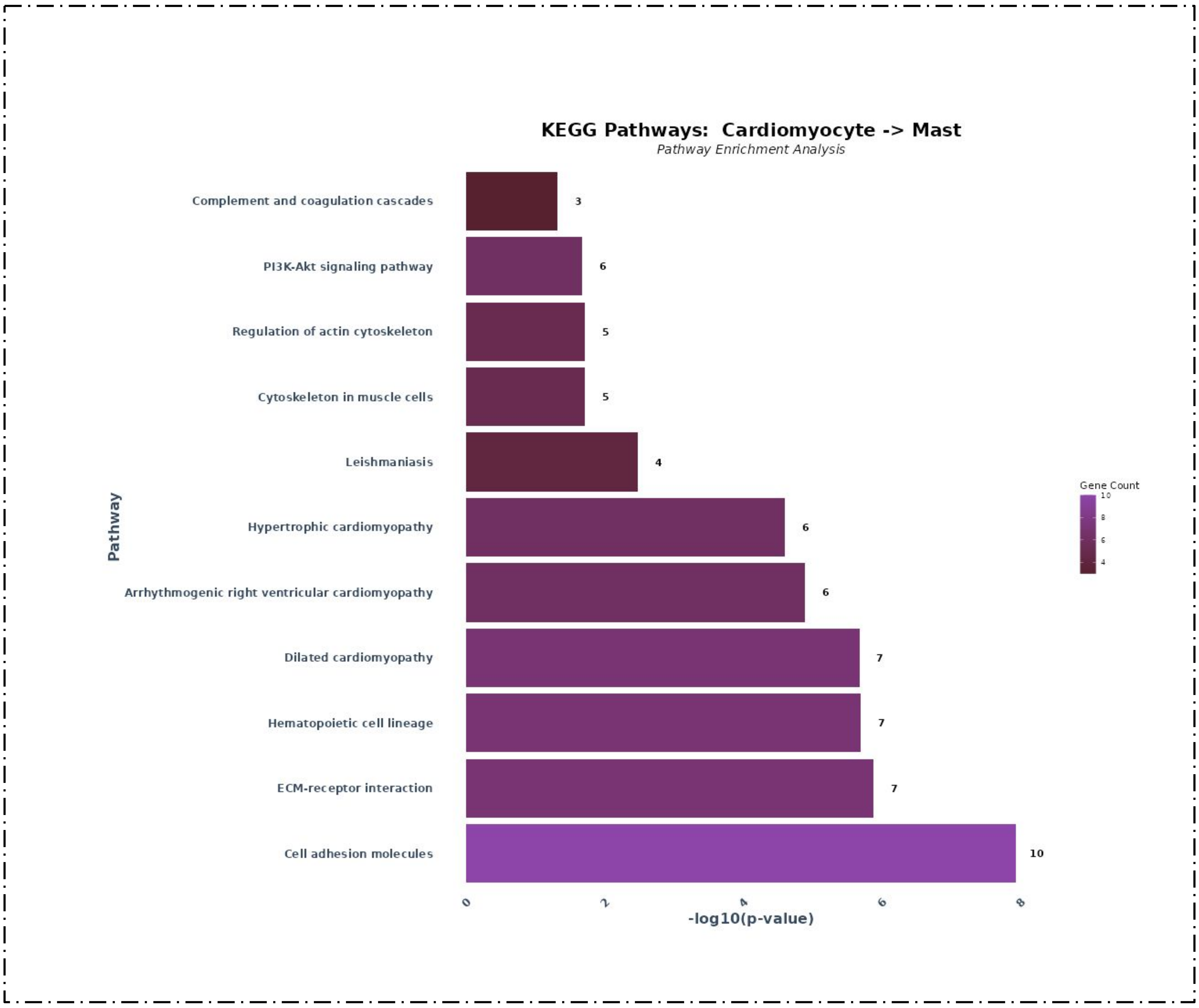


**Supplementary Figure 6.** After selecting condition and source cluster of interest, one or more target cluster are chosen. Based on the receptors present in target cluster that interact with ligands of source cluster, the KEGG analysis is done.

### Panel - Condition specific KEGG analysis

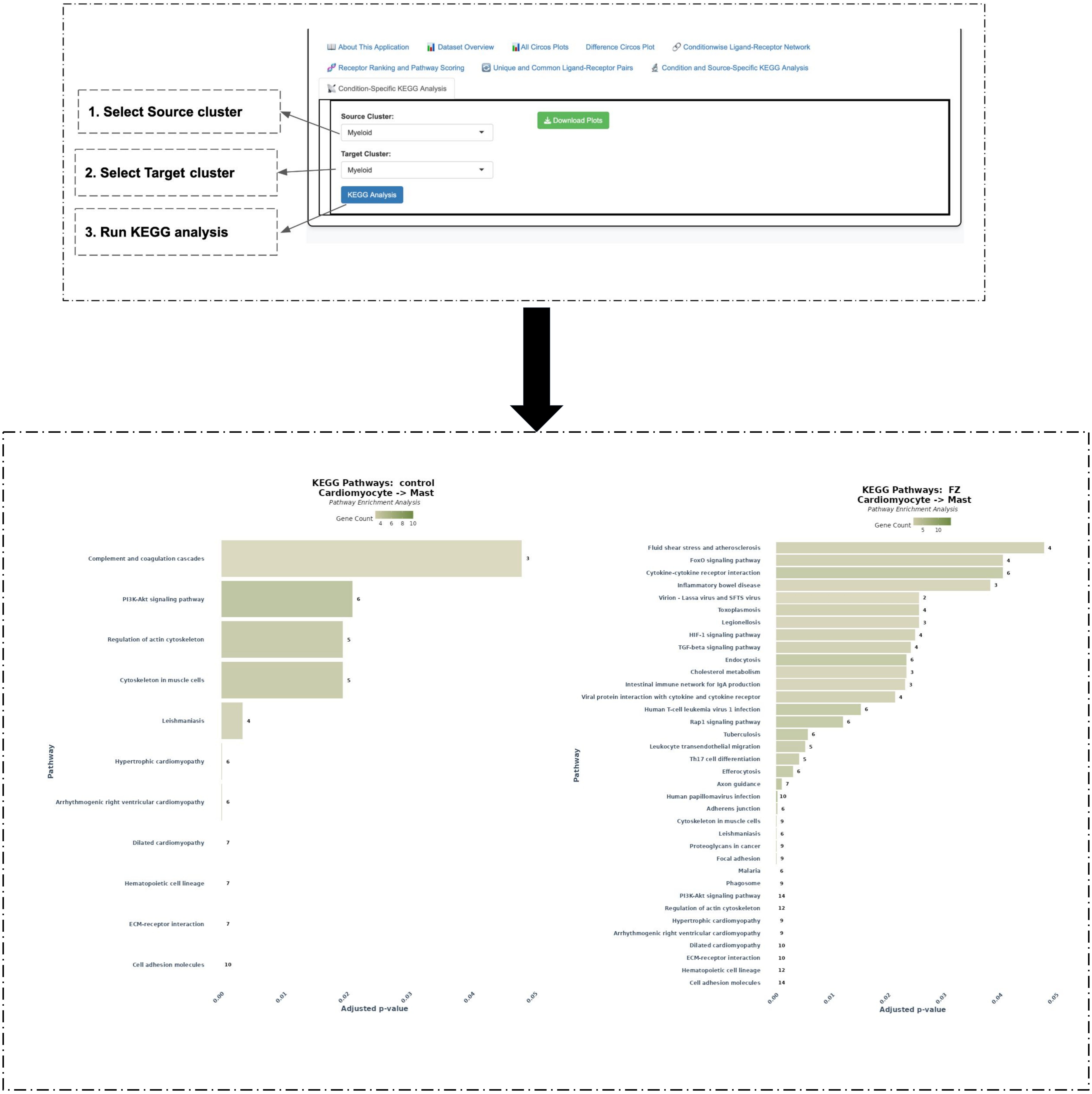


**Supplementary Figure 7.** After selecting source and target cluster of interest, and running the KEGG analysis, we get all the pathways associated with the receptor in target cluster that interact with ligands of source cluster, for every condition.
